# *TFOFinder*: Python program for identifying purine-only double-stranded stretches in the predicted secondary structure(s) of RNA targets

**DOI:** 10.1101/2023.04.26.538412

**Authors:** Atara Neugroschl, Irina E. Catrina

## Abstract

Nucleic acid probes are valuable tools in biology and chemistry and are indispensable for PCR amplification of DNA, RNA quantification and visualization, and downregulation of gene expression. Recently, triplex forming oligonucleotides (TFO) have received increased attention due to their improved selectivity and sensitivity in recognizing purine-rich double-stranded RNA regions at physiological pH by incorporating backbone and base modifications. For example, triplex forming peptide nucleic acid (PNA) oligomers have been used for imaging structured RNA in cells and inhibiting influenza A replication. Although a handful of programs are available to identify triplex target sites (TTS) in DNA, none are available that find such regions in structured RNAs. Here, we describe *TFOFinder*, a Python program that facilitates the identification of intramolecular purine-only RNA duplexes that are amenable to forming parallel triple helices (pyrimidine/purine/pyrimidine). We performed genome- and transcriptome-wide analyses of TTS in *Drosophila melanogaster* and found that only 0.3% (123) of total unique transcripts (35,642) show the potential of forming 12-purine long triplex forming sites that contain at least one guanine. Using minimization algorithms, we predicted the secondary structure(s) of these transcripts, and using *TFOFinder*, we found that 97 (79%) of the identified 123 transcripts are predicted to fold to form at least one TTS for parallel triple helix formation. The number of transcripts with potential purine TTS increases when the strict search conditions are relaxed by decreasing the length of the probes or by allowing up to two pyrimidine inversions or 1-nucleotide bulge in the target site. These results are encouraging for the use of modified triplex forming probes for live imaging of endogenous structured RNA targets, such as pre-miRNAs, and inhibition of target-specific translation and viral replication.

## Introduction

In 1957, four years after Watson and Crick published the structure of double-stranded DNA, Felsenfeld, Davies, and Rich reported the characterization of poy(A)/poly(U) triple helix formation [1]. Since then, it has been revealed that DNA and RNA triple helices have important biological roles in catalysis, regulation of gene expression, and RNA protection from degradation (reviewed in [2]).

When meeting certain requirements, nucleic acids can form triple or quadruple helices. The latter is formed by G-rich sequences and recent studies revealed quadruplex selective recognition for *in vivo* analysis of human telomeric G-quadruplex formation [3]. Natural intramolecular triple helices form for nucleic acid sequences rich in consecutive purine (R) and pyrimidine (Y) stretches and were proposed to control gene expression by inhibiting transcription or preventing the binding of other factors [4]. Intermolecular triple helices are promising tools for artificial control of gene expression and as therapeutics approaches to address various human diseases [5–7], which form when a third strand interacts with a canonical duplex via Hoogsteen base pairs (bp) (Fig 1; reviewed in [2]). The third strand can bind to the major or minor groove of a duplex; however, the minor groove triplex is unstable. In addition, depending on sequence composition, the third strand can bind in a parallel or antiparallel orientation to form Y⦁R:Y and R⦁R:Y triple helices, respectively. Where “⦁” and “:” denote Hoogsteen and Watson-Crick hydrogen-bonding, respectively. Triplex-forming oligonucleotides (TFO) can have a DNA or RNA backbone, and when they have a length of at least 10-12 nucleotides (nt), triplex formation can be characterized with common assays, such as native gel electrophoresis [8]. With an unmodified TFO (DNA or RNA), triplex formation involves the interaction between three strands all with a negatively charged frame, which leads to electrostatic repulsion and a very slow association of the third strand. However, once formed, parallel triple helices are very stable with half-lives of days. The peptide nucleic acid (PNA) backbone modification has been employed to eliminate this unfavorable interaction, which resulted in high TFO binding specificity and sensitivity, and in a greater mismatch discrimination as compared to using DNA or RNA TFO [9–11]. Triplex formation can further be favored and stabilized by employing base modifications [11–17].

**Fig 1.**
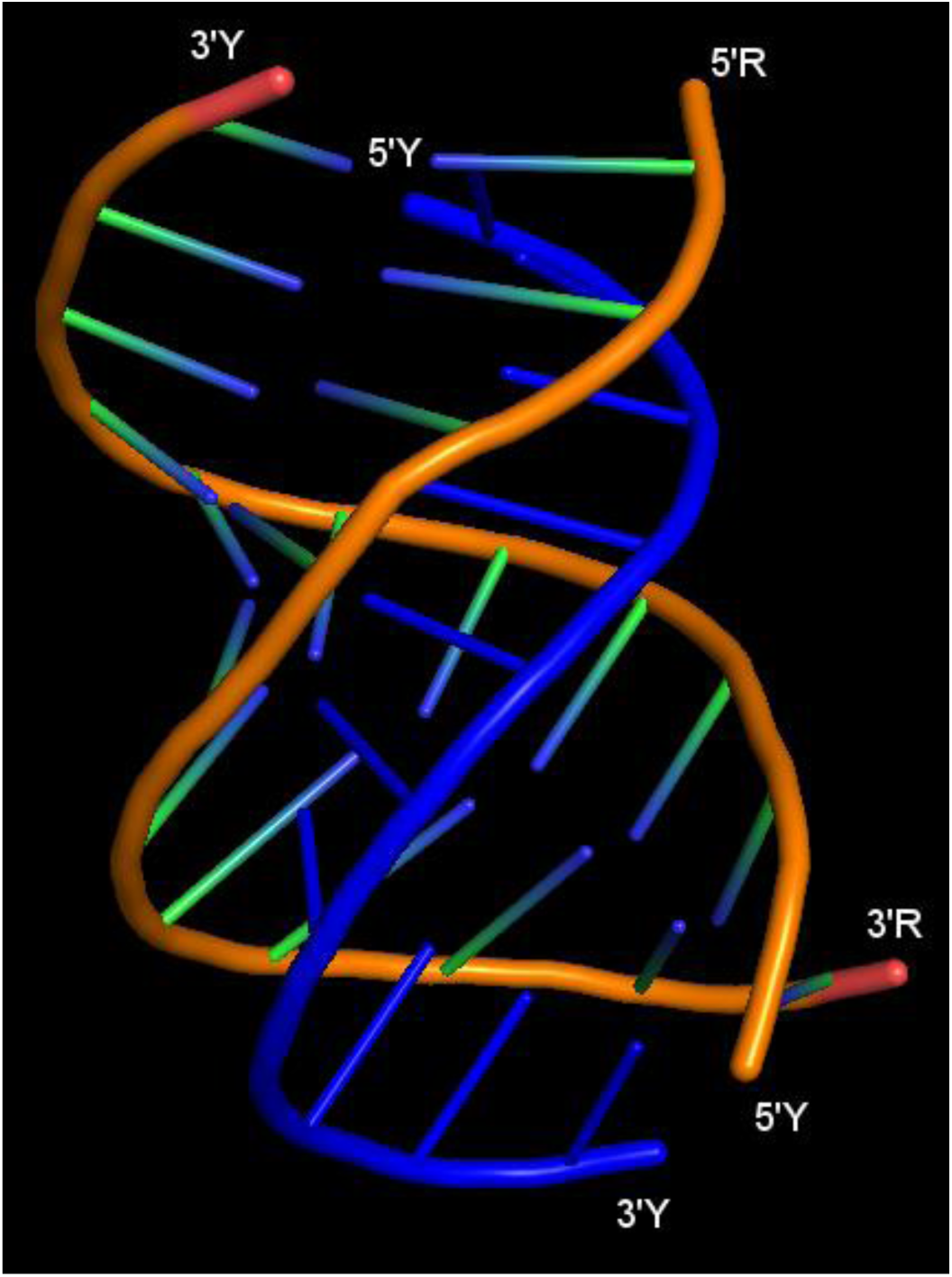
Structure of an 11-nt long intramolecular Y⦁R:Y triple helix as determined using X-ray crystallography. The strands forming the R:Y Watson-Crick duplex are shown in orange, and the triple helix forming Y strand is shown in blue. R = purine, Y = pyrimidine, “⦁” = Hoogsteen H-bonding, “:” = Watson-Crick H-bonding. Structure adapted from PDB ID: 6SVS [18].

Endogenous DNA and RNA triple helices have important biological roles; RNA splicing (RNA⦁RNA:RNA) and telomere synthesis (RNA⦁DNA:DNA) involve formation of short triple helices [19, 20]. In the first example, the backbone phosphates bind metal ions needed for splicing, and in the second example, the triplex formation is required for catalysis. Triple helices are also involved in gene expression regulation by mediating ligand binding for metabolite-sensing riboswitches in bacteria and facilitate RNA protection from degradation [21–26]. Exogenous RNA triple helices have great potential for application in imaging of endogenous RNAs, target-specific inhibition of translation, and inhibition of pre-miRNA processing.

The use of unmodified TFOs (DNA or RNA) is limited in general by the formation of intermolecular structure or motifs (I-motif and G-quadruplex) or duple-formation with single-stranded regions of target and non-target RNAs. Important advances have been made in identifying backbone and base modifications to enhance TFO selectivity. These are greatly expanding TFO applications to imaging and studies of regulation of gene expression.PNA⦁RNA:RNA triple helix formation was shown to efficiently inhibit viral replication of influenza A (IAV) [27].

Although TFOs show great promise for applications in biology and medicine, there are also a few aspects that still need to be improved:

1. Cellular, cytoplasmic, and nuclear, delivery of TFO; efficient oligonucleotide delivery is currently achieved using various delivery agents (*e.g.*, polyamines, liposomes) and/or electroporation methods, depending on the specimen and delivery site of interest. Recently, modified oligomers showed superior cellular uptake without the use of carriers [27, 28].
2. Solubility of PNA-derived TFO; exchanging the negatively charged phosphate diester for an uncharged peptide backbone coupled with the hydrophobicity of the nitrogenous bases can yield PNA oligomers with reduced water solubility. This is addressed by the addition of up to three positively charged amino acid residues, usually lysine, at the N- or C-terminus of the TFO.
3. TFO design for RNA targets; TFO design for double-stranded DNA targets is straightforward, one only needs to search the target DNA sequence for purine stretches with the length of interest. The *Triplexator* application was reported to predict short (< 30-bp) double-stranded DNA binding sites for a given RNA sequence [29]. *LongTarget* finds longer DNA TTS, the *Triplex Domain Finder* application detects DNA-binding domains in long non-coding RNAs, and the *Triplex* from the *R/Bioconductor* suite predicts the formation of eight types of intramolecular triplexes within a given nucleic acid sequence [30–32]. However, to our knowledge, there are no applications that facilitate the design of TFO for structured RNA targets containing R:Y duplex regions, which can form intermolecular triplexes.

Here, we describe *TFOFinder*, an open-source Python program to design parallel pyrimidine TFO recognizing purine-only double-stranded regions in any RNA target of interest (Y⦁R:Y). We used *RNAMotif* and *TFOFinder* to determine the prevalence of potential DNA, and RNA target sites in the *Drosophila melanogaster* genome (*version 6.48*) and transcriptome (*version 6.38*), respectively [33]. *RNAMotif* uses descriptor files to search for a user-defined primary or secondary structure “motif” within a given file containing one or more sequences in the FASTA format [33].

DNA and RNA triple helices have been extensively analyzed via optical melting experiments, circular dichroism, FRET, and other techniques. Of particular interest are RNA⦁DNA:DNA and RNA⦁RNA:RNA triple helices, as they have essential biological roles, such as telomere synthesis where they ensure proper pseudoknot folding, catalysis without direct association with the active site, and recruiting divalent metal ions for splicing (reviewed in [2]). Efficient triple helix formation with a TFO containing an unmodified DNA/RNA backbone requires at least 10-bp long purine rich TTS and a mildly acidic pH to protonate cytosines such that they can participate in Hoogsteen base pairing. TTS hairpin models with purine-rich stems and random loop sequence are commonly used to analyzed TFO properties in solution (Fig 2).

**Fig 2.**
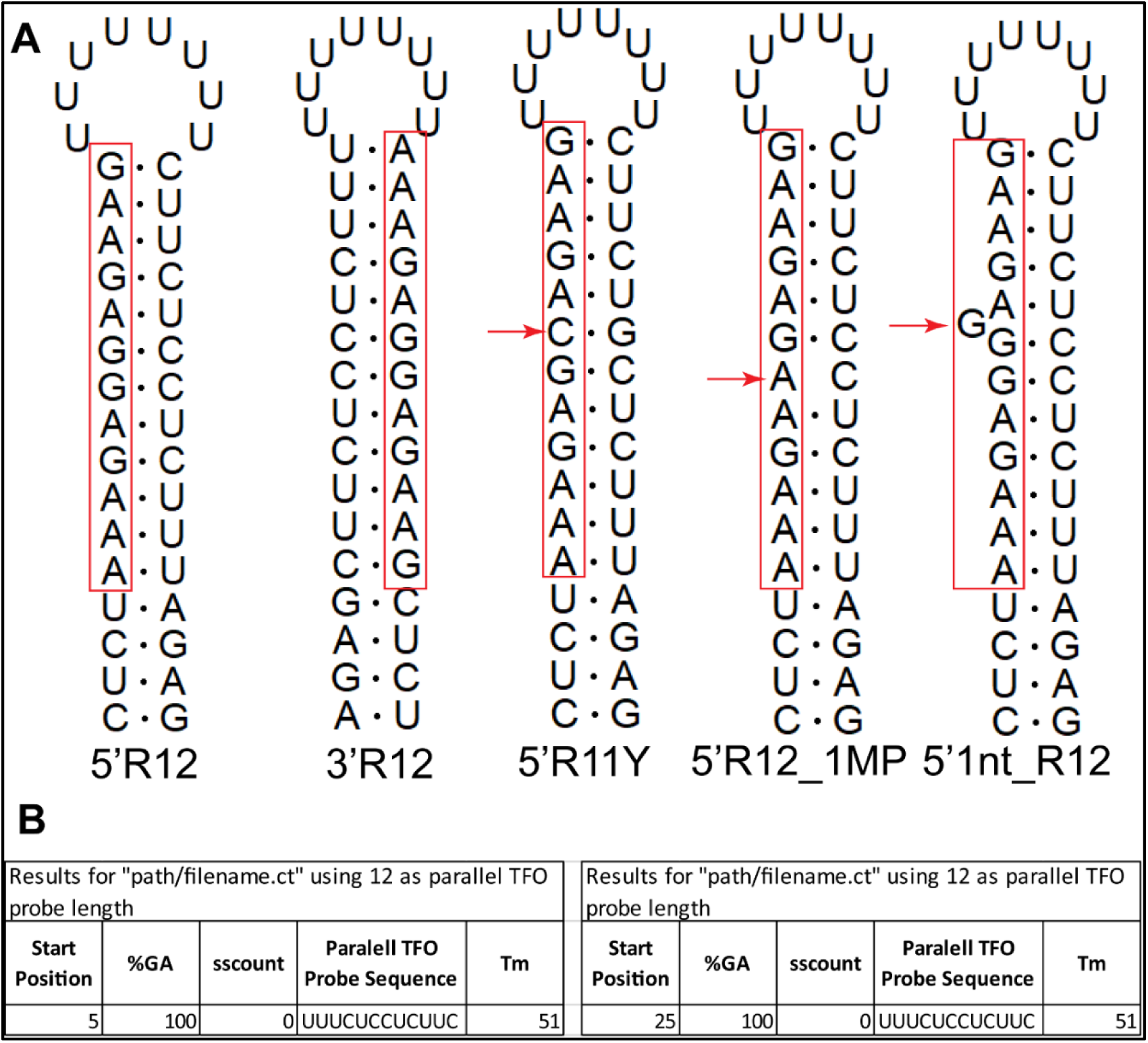
Model RNA hairpins illustrating examples of ideal and interrupted 12-bp long TTS. (A) The purine stretch can be positioned on the 5’ (5’R12) or 3’ (3’R12) side of the hairpin duplex, and these two TTS are readily identified by *TFOFinder*. The remaining three TTS are not reported by *TFOFinder* and can only form stable triplexes with a modified TFO. 5’R12Y = the purine region is positioned on the 5’ side of the duplex and it is interrupted by a pyrimidine inversion (red arrow). 5’R12_1MP = the purine region is positioned on the 5’ side of the duplex and it is interrupted by a mispair (red arrow). 5’1nt_R12 = the purine region is positioned on the 5’ side of the duplex and it is interrupted by a 1-nt bulge (red arrow). (B) The *TFOFinder* output for the first two TTS RNA hairpin examples, 5’R12 and 3’R12.

We show that our program facilitates the identification within any RNA target of duplex regions amenable to forming a parallel Y⦁R:Y triplex, and the design of the corresponding short TFO probes (4-30-nt). These TFO probes can be used for specific inhibition of translation and imaging of structured RNAs containing purine-rich sequences in non-denaturing conditions.

## Results and Discussion

TFO probes have already found important applications in imaging of cellular RNAs and nucleic acid function modulation and assays [27, 34–37]. Here, we explored the feasibility of extending the application and versatility of TFOs by performing a transcriptome- and genome-wide analysis in *D. melanogaster* to identify all RNA and DNA stretches that are amenable to triple helix formation. Moreover, we tested our program by designing TFO probes for a previously reported RNA target, the vRNA8 of influenza A, which encodes two essential viral proteins, NEP and NS1 [27, 38, 39]. TFO probes designed using *TFOFinder* are promising tools for *in vivo* imaging of structured RNA targets (*e.g.*, pre-miRNAs), determining *in vivo* folding of endogenous RNA targets, target-specific inhibition of translation, and others.

To identify continuous single-stranded stretches of 12 purines, we searched the fruit fly transcriptome using *RNAMotif*, a program that finds user-defined sequences or potential structural motifs in a given nucleic acid target sequence without information about the target’s secondary structure [33]. We counted adenine (A)-only stretches separately from guanine (G)-containing ones and identified all hits corresponding to unique transcripts. We then searched the sequence of the transcripts containing these hits for a complementary match, or a match containing G-U wobble pair(s), or with one mispair.

### Drosophila melanogaster genome survey

Both strands of the DNA genome were searched for R12 stretches [containing as least one G], which were identified and counted for defined DNA regions (Table 1). These stretches were found in more than 50% of targets for intronic, mRNA, and gene sequences. The largest number of hits were obtained for intronic regions (437,487), mapped to 23.72% of total unique intronic targets. tRNAs and miRNAs contained the least number of R12 sequences, mapped to only 0.96% (3) and 2.68% (20) of total tRNA and miRNA unique targets. However, not all R12 hits listed in Table 1 are unique, as the exon, UTRs, gene, and mRNA sequences present significant overlap.

**Table 1.**
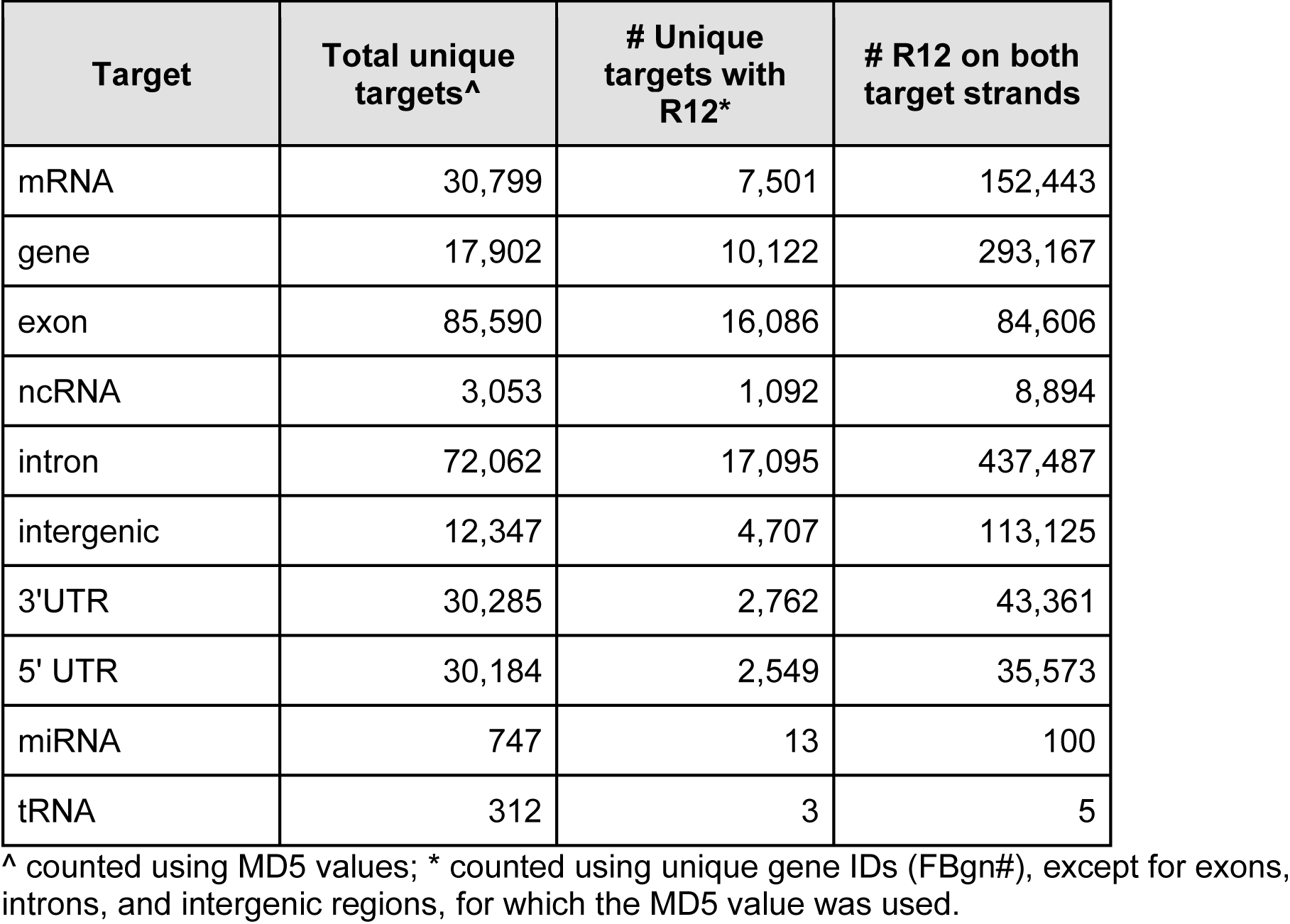
Results for the *D. melanogaster* genome (*v. 6.48*) for R12.

### Drosophila melanogaster transcriptome survey

While triple helix formation with a DNA/RNA TFO requires the presence of a continuous stretch of purines in the target, it has been shown that triplexes can be formed with TTS containing one or two pyrimidine inversions (Fig 2, 5’R11Y) when a modified TFO is employed. Therefore, we determined whether allowing pyrimidine inversions would significantly increase the number of transcript hits. We analyzed the full *D. melanogaster* transcriptome by beginning with a strict search (R12 - all purines and not all As), which we gradually relaxed to allow for G**-** U pairing, or one mispair (Fig 2, 5’R12_1MP), or up to two pyrimidine inversions in accordance with previously reported triple helix formation rules and restrictions (Table 2, Fig 3) [12, 14, 40-42]. For the strictest search, for R12 sequences, we identified all 12-nt stretches of purines that had at least one G and found that 123 unique transcripts (0.3% of the total 35,642 transcripts) also contained at least one corresponding complementary sequence needed to form an R12 TTS, which were encoded within 54 unique genes (0.3% of the total 17,787 genes; Table 2, R12; S1 Table). When G-U paring was allowed, we identified 1,506 (4.2%) unique transcripts mapped to 620 (3.5%) unique genes containing complementary sequences with the potential of forming 12-bp long purine duplexes (Table 2, R12_GU). When one mispair was allowed, there were 811 (2.3%) unique transcripts mapped to 351 (2.0%) unique genes containing complementary sequences with the potential of forming 12-bp long purine duplexes (Table 2, R12_1MP). When we relaxed the conditions to allow for one pyrimidine inversion (Table 2, R11Y), and eliminated the requirement for a G, 391 (1.1%) unique transcripts were identified, corresponding to 178 (1.0%) unique genes (Table 2, R11Y). Finally, we also allowed for two pyrimidine inversions (R10Y2). First, we restricted the position of the inversions to the middle of the TTS, and not consecutive. With these search restrictions we found 317 (0.9%) unique transcripts, corresponding to 138 (0.8%) unique genes (Table 2, R10Y2 strict). Second, we relaxed the R10Y2 search to allow the two internal pyrimidine inversions to be consecutive and/or terminal. Under these conditions, we discovered 606 (1.7%) unique transcripts mapped to 269 (1.5%) unique genes (Table 2, R10Y2 relaxed).

**Fig 3.**
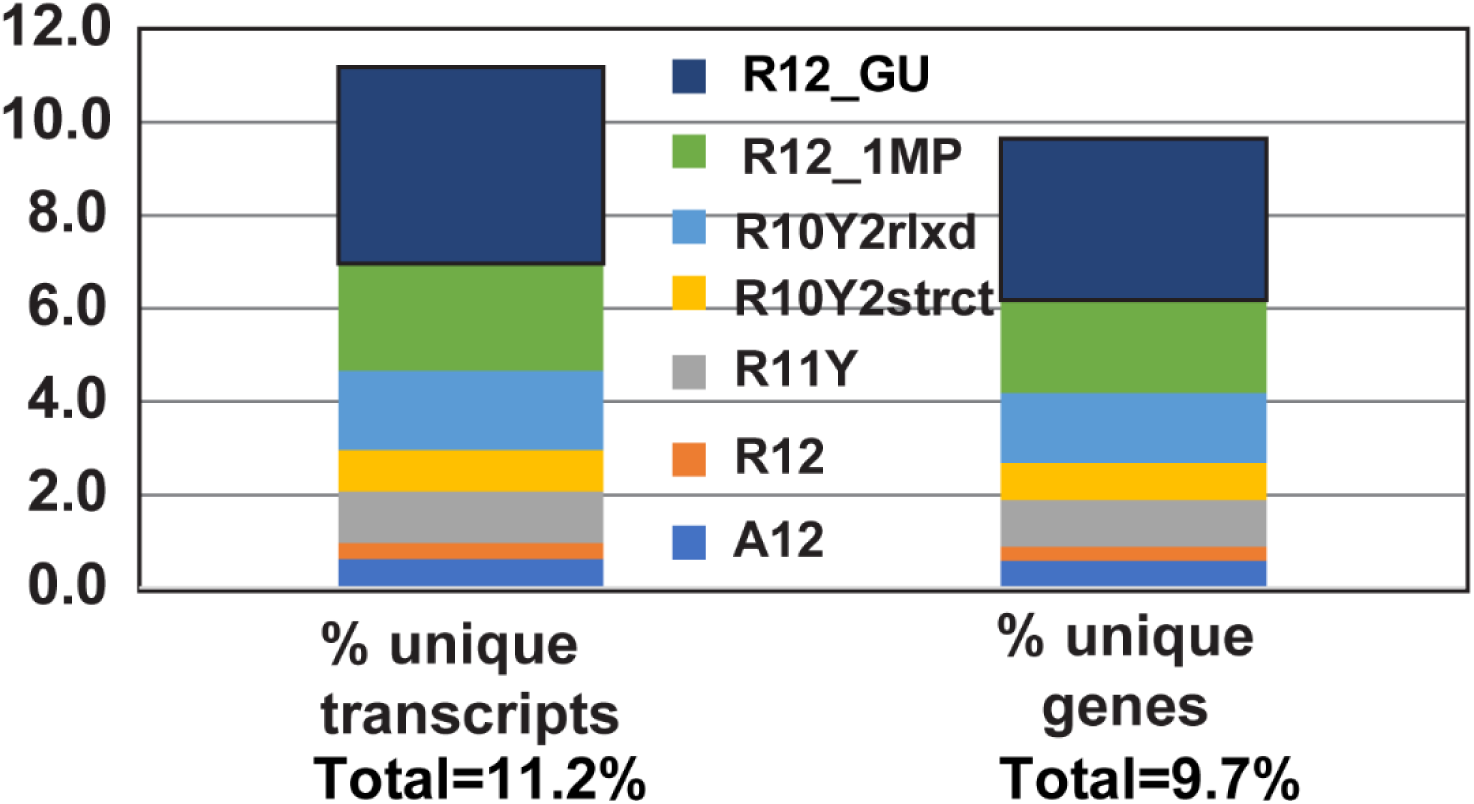
The number of unique transcripts and corresponding genes containing the indicated TTS types, as obtained from the transcriptome (*v. 6.38*) analysis. A12 = 12 consecutive adenines; R12 = 12 consecutive purines, containing at least one guanine; R12_1MP = a duplex of R12 with 1 mispair/mismatch; R11Y = 12 consecutive nucleotides composed of eleven purines (A11 or AiGj with i+j = 11) and one pyrimidine; R10Y2 = 12 consecutive nucleotides composed of ten purines (A10 or AiGj with i+j = 10) and two pyrimidines; strict (strct) = two Ys not next to each other and not at the ends; relaxed (rlxd) = two Ys anywhere. Total number of transcripts = 35,642; Total number of genes = 17,878.

**Table 2:** Results for the survey of *D. melanogaster* transcriptome (*v. 6.38*) for the indicated purine-rich sequences.

| Sequence | Total Single-stranded hits | Single-stranded: # unique transcripts | Single-stranded: # unique genes | Total Double-stranded hits | Double-stranded: # unique transcripts | Double-stranded: # unique genes |
| --- | --- | --- | --- | --- | --- | --- |
| **A12** | 8,707 | 2,300 (6.5%) | 1,031 (5.8%) | 2,453 | 217 (0.6%) | 105 (0.6%) |
| **R12** | 128,139 | 16,689 (46.8%) | 7,076 (39.6%) | 494 | 123 (0.3%) | 54 (0.3%) |
| **R12_GU** | n/a | n/a | n/a | 31,213 | 1,506 (4.2%) | 620 (3.5%) |
| **R12_1MP** | n/a | n/a | n/a | 5,046 | 811 (2.3%) | 351 (2.0%) |
| **R11Y** | 588,205 | 30,793 (86.4%) | 14,601 (81.7%) | 813 | 391 (1.1%) | 178 (1.0%) |
| **R10Y2 strict** | 1,402,460 | 33,935 (95.2%) | 16,733 (93.6%) | 438 | 317 (0.9%) | 138 (0.8%) |
| **R10Y2 relaxed** | 2,663,079 | 34,259 (96.1%) | 16,993 (95.0%) | 890 | 606 (1.7%) | 269 (1.5%) |

A12 = 12 consecutive adenines; R12 = 12 consecutive purines, containing at least one guanine; R12\_1MP = a duplex of R12 with 1 mispair/mismatch; R11Y = 12 consecutive nucleotides composed of eleven purines (A11 or AiGj with i+j = 11) and one pyrimidine; R10Y2 = 12 consecutive nucleotides composed of ten purines (A10 or AiGj with i+j = 10) and two pyrimidines; strict = the two Ys not next to each other and not at the ends; relaxed = two Ys anywhere. Total number of transcripts = 35,642; Total number of genes = 17,878.

Using the PANTHER Classification System, we performed a gene ontology enrichment analysis for molecular functions and biological processes for the 54 unique genes from which the 123 transcripts with potential TTS are expressed [43]. This analysis identified 13 (28.3%) genes with binding as molecular function, 14 (30.4%) and 13 (28.3%) genes involved in biological regulation and metabolic processes, respectively. However, 50% or more of these genes were not assigned to any PANTHER category.

### TFOFinder

The *TFOFinder* program, to our knowledge, is the first to search within the predicted secondary structure(s) of an RNA target of interest for double-stranded fragments of a user-defined length (4-30-nt) that are composed of consecutive purines (*e.g.*, R12, R12_GU, and A12). The *TFOFinder*’s flow chart shows the main steps of the program (Fig 4). The program identifies purine-only regions that are double-stranded and can include G-U wobble pairs, within the RNA target secondary structure(s) predicted via energy minimization algorithms (*e.g.*, *mfold*, *RNAstructure*). Moreover, the program disregards any hits that present a bulge loop on either side of the double strand. In other words, both strands are composed of only consecutively paired nucleotides. The input file is the “ct” output file from the *mfold*, *RNAstructure*, or *RNAFold* program [44], which is a common text file format for writing nucleic acid secondary structure.

**Fig 4.**
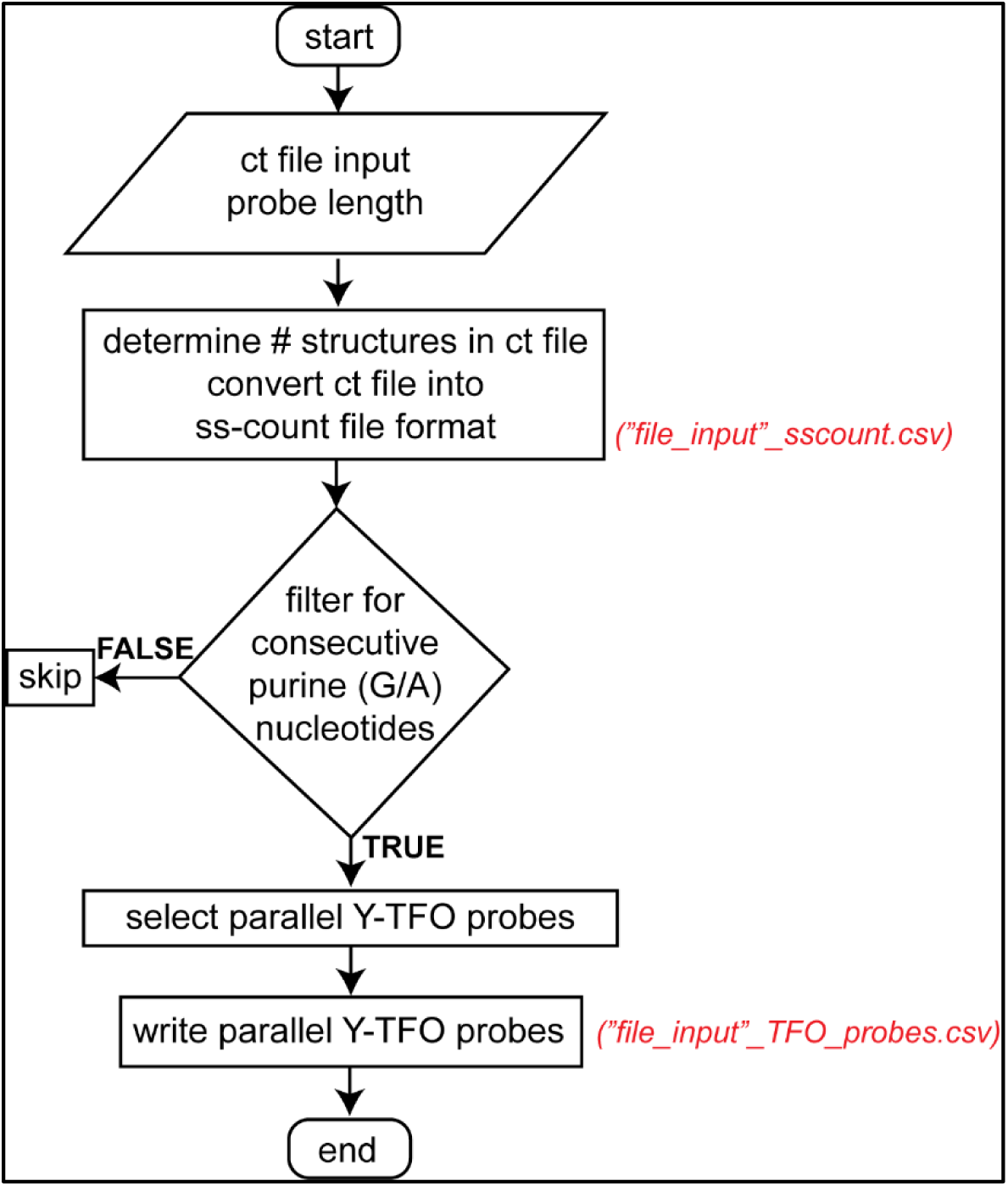
***TFOFinder* program flowchart.**

The *TFOFinder* output file lists the most 5’ number for the position of the duplex regions identified in the RNA target structure, parallel pyrimidine probe sequence for a user-defined length between 4 and 30 nucleotides and melting temperature for an intermolecular duplex between the TFO RNA and the corresponding complementary RNA sequence (Fig 2B). A target region is identified as a hit if it is predicted to form a R:Y (including G-U pairs) uninterrupted duplex when considering base pairing in all predicted secondary structures for the RNA target of interest (*i.e.*, MFE and SOs). When SO structures are included in the “ct” input file, a nucleotide will be considered as double-stranded if it has a corresponding pairing nucleotide in at least one of the structures.

### TFOs for *D. melanogaster* RNA targets

We previously found that it is beneficial to take into consideration predicted suboptimal structures when designing molecular beacon probes for live cell imaging [45]. However, computational time significantly increases when applying minimization algorithms to folding long RNA targets (>11,000-nt), and a dynamic programming algorithm has been shown to not only produce the MFE structure much faster, but also with improved accuracy for long RNA targets [46]. We used *mfold*, *RNAstructure*, and *LinearFold* to predict the secondary structure of the 123 unique transcripts identified in our *RNAMotif* search, and we analyzed the distribution of the 494 total TTS hits (Table 2, R12 – total double-stranded hits) between the MFE and SO structures (Table 3). We found that 21% (26) of the targets identified using *RNAMotif* did not present a predicted 12-bp duplex amenable to forming an Y⦁R:Y triplex within their secondary structure(s), while for 19% (23) RNA transcripts the SO structures presented TTS, but the MFE structure did not.

**Table 3:** Distribution of the 494 TTS hits identified in 123 unique transcripts between the MFE and SO structures, which were predicted using minimization algorithms (*mfold*, *RNAstructure*, *LinearFold*)

| MFE TTS | SOs TTS | # Transcripts |
| --- | --- | --- |
| none | none | 14 |
| none | N/A | 12 |
| x | x | 27 |
| none | > 1 | 23 |
| x | y > x | 33 |
| > 1 | N/A | 14 |
| **Total # transcripts** |  | **123** |

x, y > 0 are number of transcripts with R12 TTS in the MFE and/or SO structures.

For TFO targeting to work as intended, probe specificity and sensitivity are essential characteristics. We analyzed the specificity of the TFO probes identified for the 123 transcripts by analyzing all TTS sequences and, of the 4,095 possible unique TTS R12 sequences, 50 were found in the 494 total R12-double-stranded hits (Table 4), with two R12 sequences composed of consecutive “GA” or “AG” representing 45% of total hits (223 of 494; Table 4), and contained within 13 unique transcripts mapped to three unique genes (*eag*, *RSG7*, and *CG42260*). Further analysis of these TTS sequences showed that 48% (21 of 50; Table 4) of the identified TTS were unique sequence hits and were mapped to 12 unique transcripts encoded within 12 unique genes.

**Table 4:** Distribution of the *TFOFinder* identified TTS sequences.

| # TTS occurrences | # unique TTS | # unique transcripts | # unique genes | # hits |
| --- | --- | --- | --- | --- |
| ≥ 100 | 2 | 13 | 3 | 223 |
| ≥ 10 | 11 | 93 | 38 | 192 |
| ≥ 2 | 16 | 39 | 16 | 58 |
| 1 | 21 | 12 | 12 | 21 |
| **Total number of TTS hits** |  |  |  | **494** |

We sorted the 123 transcripts according to their length, and the first two transcripts were two noncoding RNAs (*CR44598-RA*, 486-nt and *CR44619-RA*, 1,023-nt). For the first one, one TTS was identified in all target structures (MFE and 13 SO structures; 5’ location = 246, sscount = 0.68) (Fig 5A), while for the second one, five non-redundant TTS were identified in the suboptimal structures, but none were found in the MFE structure. For example, the TTS mapped between 880-892 was present in five of 19 total SO structures (Fig 5B), while in the MFE structure it presented a mispair (MP) and a 1-nt bulge (Fig 5B, arrows). In addition, the sscount value for a TFO probe should be as close to zero as possible, as the sscount being equal to zero means that all nucleotides are base-paired.

**Fig 5.**
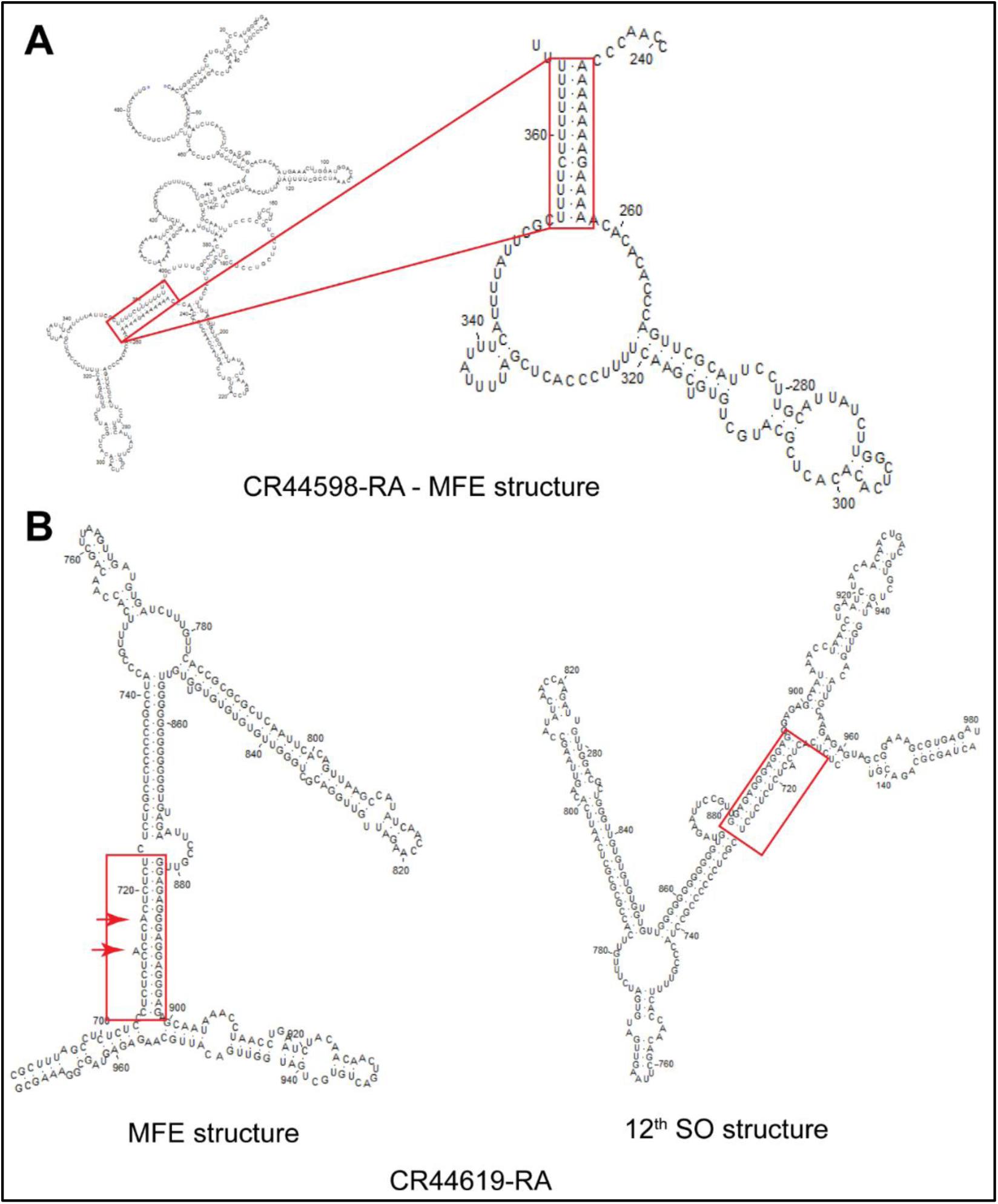
Secondary structures for two ncRNAs, predicted with *mfold*. (A) Full MFE structure (*mfold*) of the shortest transcript (*CR44598-RA*) identified to contain one TTS, which is highlighted in the red box and also shown magnified (right). (B) A longer ncRNA (*CR44619-RA*, 1,023-nt) containing several TTS, one highlighted in the red box for the of the 12^th^ SO structure (*mfold*; right) and containing a mispair and 1-nt bulge in the MFE structure (left, red arrows).

### TFOs for *Influenza A* vRNA8 target

Using *TFOFinder*, we explored a previously reported RNA target that was shown to form PNA⦁RNA:RNA triplexes *in vivo* [27]. Partially complementary sequences at the 5’ and 3’ end of all eight vRNAs of IAV make up a conserved panhandle motif that acts as a viral promoter for transcription and replication. However, this motif contains at least one bulge and therefore it does not fit the ideal requirements for parallel Y**·**R:Y triple helix formation and requires TFO modification to form a triplex. The panhandle region of vRNA8 was identified as a TFO-target and it was reported that a modified PNA TFO efficiently inhibits IAV replication [27]. Using the Clustal 1.2.4 webserver [47], we performed a sequence alignment of 15 vRNA8 Viet Nam strains and found that the reported TTS was not conserved among these sequences, which means that the identified TFO would work only for the HM006763A strain.

Therefore, using *TFOFinder*, we searched for additional TTS in the same 15 vRNA8 IAV Viet Nam sequences, and compared our results with the experimentally probed secondary structure of the target vRNA [38]. When including in the search the MFE and SO structures, we identified three conserved regions, two of which are highlighted in Fig 6 (green boxes, TTS positioned at 365 and 218) (Table 5). This means that the MFE structure did not present an ideal TTS, but each purine contained in the identified TTS were double-stranded in at least one of the SO structures. The first and third TTS (Table 5: TTS positioned at 365 and 804) do not appear to be good candidates to form a triplex because the former is part of a multibranch loop and the latter includes an internal loop [38]. However, the reported structure was determined using solution assays and it is possible that the *in vitro* structure may differ from the *in vivo* folding of the RNA target, although one would expect the *in vivo* folding to be less structured [48]. The second TTS (Table 5, TTS positioned at 218) may be a viable alternative and is conserved in all strains, but it is shorter than the recommended minimum length (8 vs. 10-nt), which may compromise the sensitivity and specificity of the assay for the targeted TTS. To assess the specificity of this probe, using *RNAMotif*, we performed for the IAV TTS-218 similar searches as described for the *D. melanogaster* transcriptome for both *D. melanogaster* (*v. 6.38*) and *H. sapiens* (May 23^rd^, 2018) transcriptomes (Table 6). We found that in *D. melanogaster*, only 0.04% of transcripts had the potential to form the double-stranded IAV TTS-218, while in *H. sapiens* this percentage increased to 2.58%, which was still small. However, a longer TTS would make a more attractive region to design modified TFOs for functional inhibition.

**Fig 6.**
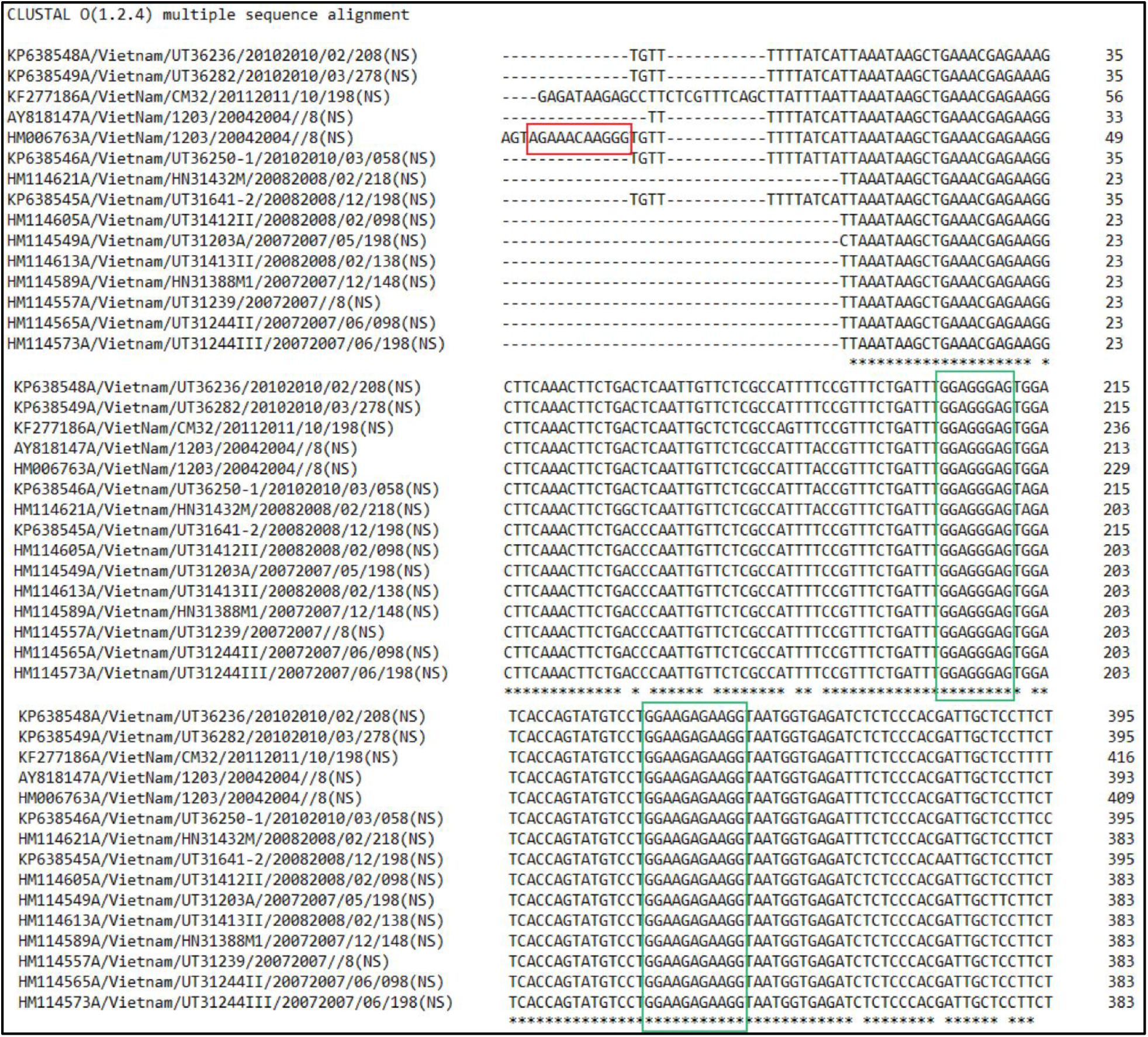
Alignment of 15 vRNA8 IAV sequences (*Clustal 1.2.4* webserver) The red box highlights the panhandle TTS experimentally targeted for inhibiting influenza A replication. The green boxes highlight two conserved TTS identified using *TFOFinder*.

**Table 5:**
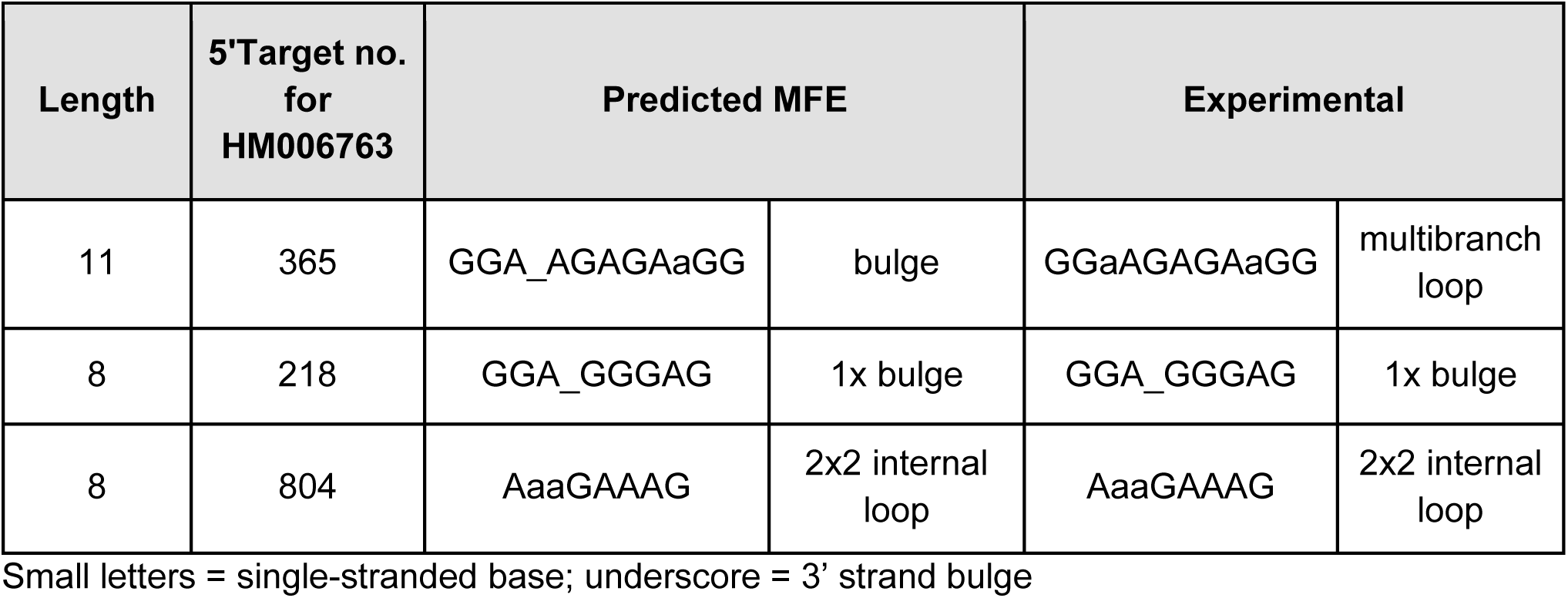
*TFOFinder* results for vRNA8 IAV Viet Nam strain HM006763.

**Table 6:**
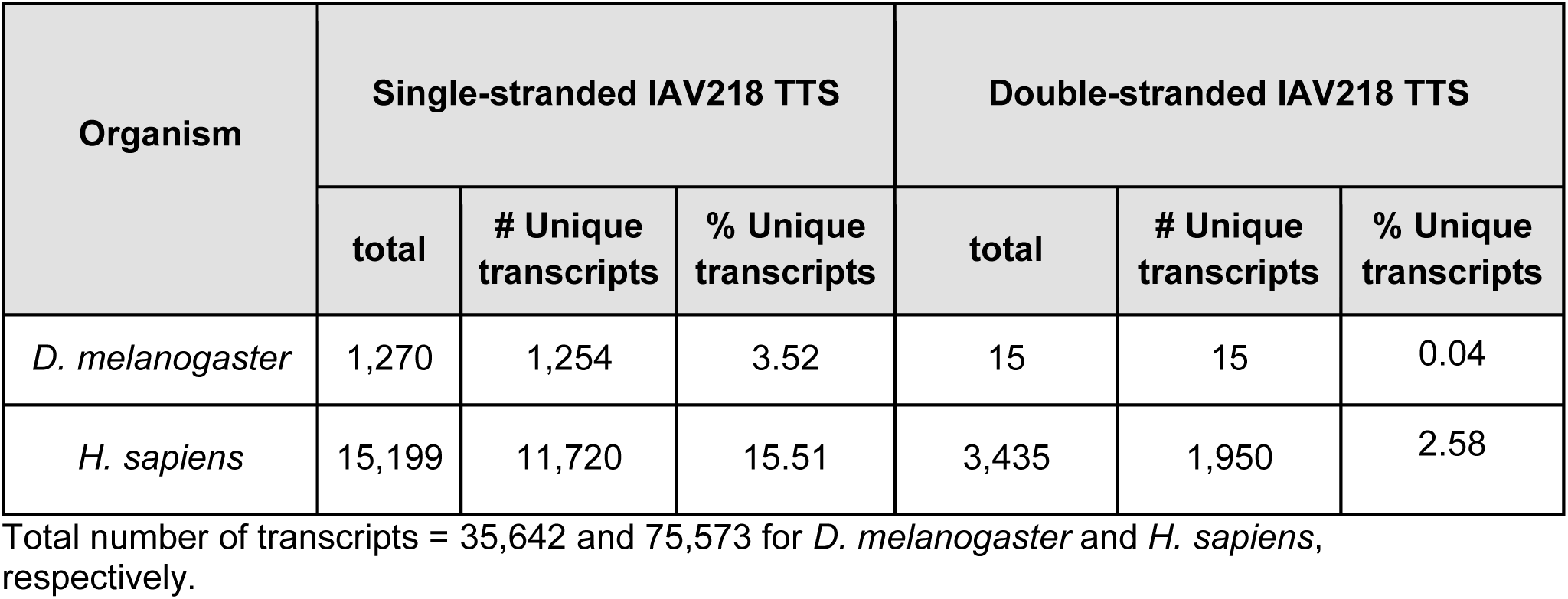
*RNAMotif* results for IAV TTS-218 prevalence in *D. melanogaster* and *H. sapiens* transcriptome.

## Conclusion

*TFOFinder* is a platform independent Python program for the fast and efficient identification within any RNA structure of purine-only double-stranded regions that are predicted to form parallel triple helices of the TFO⦁RNA:RNA type. The design of target specific TFO probes is applicable to studies of *in vivo* RNA structure, RNA imaging, and RNA function regulation.

## Materials and Methods

### Target sequences

#### *D. melanogaster* transcriptome and genome

For the survey of *Drosophila melanogaster* targets, the corresponding FASTA sequences were downloaded using the *Flybase* online tools [49]. The full transcriptome *version 6.38* (02/18/2021) and genome *version 6.48* (09/26/2022) were used to perform the surveys.

#### Influenza A vRNA8

The full-length segment 8 sequences of IAV Viet Nam strain were downloaded from the NCBI Influenza Virus Resource [50]. The reverse complement of these 15 sequences, which are the vRNA sequences, were generated using *BioEdit* [51], folded using *Fold-smp* from the *RNAstructure version 6.4* [52] using the previously reported SHAPE data file and constraints (slope = 2.6 and intercept = −0.8) [38]. The resulting “ct” files, which contained information about the secondary structure of the minimum free energy (MFE) and up to 19 suboptimal (SO) structures, were used to identify TFO-target regions using a batch version of *TFOFinder*.

#### Homo sapiens refseq_rna

The FASTA sequences were downloaded from the NCBI download site last updated on May 23^rd^, 2018, using the Aspera download tool [NCBI>refseq>H_sapiens>mRNA_protein>human.X.rna.fna.gz, (X = 1, 2, 10, 11, and 12)].

### *D. melanogaster* genome survey for purine rich sequences

We searched for 12 consecutive purines, including all As [R(A)12] on both strands of the *D. melanogaster* DNA sequences downloaded in FASTA format as gene, mRNA, ncRNA, miRNA, tRNA, exon, intron, intergenic, 5’UTR, and 3’UTR. The transcriptome search described below identified single-stranded purine sequences that corresponded to the double-stranded DNA encoding each transcript. However, the transcriptome survey did not consider the intergenic and intronic parts of the DNA genome, moreover, many of the hits found in transcripts were redundant as many genes encode for several mRNA variants with overlapping sequences.

### *D. melanogaster* transcriptome survey for purine rich sequences

We identified purine-rich sequences in all *D. melanogaster* transcripts by performing one-strand searches using *RNAMotif*. We searched for single-stranded purine-only sequences composed of consecutive purines (R12) that were not all adenines (A), or contained only As (A12), or for purine-rich regions interrupted by up to two pyrimidines (R11Y and R10Y2). We next searched within the identified transcript sequences for complementary regions that can form a duplex with the already identified single-stranded hits. To confirm our results, we also performed this search on the full transcriptome, and the two searches yielded the same hits.

#### Identification of transcripts with single-stranded purine-rich stretches

Using *RNAMotif*, we scanned the transcriptome of *D. melanogaster* for stretches of 12 purines, all adenine (Table 2, A12, single-stranded), containing at least one guanine (Table2, R12_GU, single-stranded), or up to two pyrimidines (Table2, R11Y, R10Y2 strict and relaxed), single-stranded). From the *RNAMotif* output, we extracted all unique transcript IDs and downloaded their sequences in FASTA format using the *FlyBase Sequence Downloader* tool.

#### Identification of transcripts with double-stranded purine-rich stretches

Using *RNAMotif*, we then identified transcripts containing the corresponding complementary pyrimidine sequence(s) (Table 2, A12, R12, double-stranded). From the *RNAMotif* output file we extracted the transcript name, length, and genomic location, as well as their IDs using the *FlyBase Batch Download* tool. The search was then relaxed to allow for one mispair (Table 2, R12_1MP, double-stranded; Fig 2, 5’R12_1MP), or for one (Table 2, R11Y,double-stranded; Fig 2, 5’R11Y), or two pyrimidine inversions either anywhere in the 12 sequence (Table 2, R10Y2 relaxed, double-stranded) or restricted to the 10 internal positions and not consecutive (Table 2, R10Y2 strict, double-stranded). After identifying all TTS showing the potential to be double-stranded, we predicted the secondary structure(s) of the transcripts that contained them using minimization algorithms. Using *TFOFinder* we analyzed the likelihood of each TTS to be double-stranded in the predicted secondary structure(s). To find the predicted MFE secondary structure of the transcript, we used *LinearFold* for RNA targets longer than 11,000-nt, *mfold* for transcripts with up to 2,400-nt, and *RNAstructure* for the remaining sequences. In addition to the MFE structure, *RNAstructure* and *mfold* provided a various number of suboptimal structures. Using *TFOFinder*, we took into consideration the predicted secondary structure(s) to identify regions of 12 double-stranded purines.

#### Analysis of hits

Using gawk, custom Python scripts, and *Flybase* tools, we extracted the ID of the unique transcripts and the corresponding unique genes to which the hits were mapped.

### TFOFinder program

The open-source program was written in Python with a text interface, and it is freely available on GitHub (https://github.com/icatrina/tfo_rna). The input file is the “ct” format file, which is used to count the total number of structures (MFE and SO), identify consecutive purines of a user-defined length (4-30-nt) and list in the output file information for the parallel (5’ è 3’) TFO probes forming Y**·**R:Y triplexes. The output lists the 5’ start position for the identified TTS that can form a Y**·**R:Y parallel triplex, the percentage of G/A content of the RNA TTS, the parallel TFO sequence, and the melting temperature (*T*_m_) of the duplex of the RNA TFO and the corresponding complementary RNA sequence. Alternatively, the *TFOFinder* can be used via free Amazon Web Services (AWS), with AWS CloudShell, which allows for up to 1GB free persistent storage.

## Acknowledgments

We are very thankful and grateful to Dave Matthews, MD, PhD, from the University of Rochester for his continuing support with *RNAstructure* algorithms and for his invaluable help and advice on thermodynamic analysis of nucleic acid folding, programming, and more. We thank Livia V. Bayer, PhD, for critically reading this manuscript and helpful discussions. We are also grateful to the *Flybase* help team, and in particular to Josh Goodman, Julie Agapite, and Victor B. Strelets, for promptly answering our questions and writing customized scripts to meet our needs. Finally, we would like to thank to current and past members of the Catrina laboratory for their experimental work that has contributed to the planning of this computational analysis. This work was supported by a Yeshiva University start-up fund (IEC).

## Abbreviations

TTS: <u>T</u>riplex <u>T</u>arget <u>S</u>ite(s)
R: purine nitrogenous base
Y: pyrimidine nitrogenous base
bp: <u>b</u>ase <u>p</u>air(s)
TFO: <u>T</u>riplex-<u>F</u>orming <u>O</u>ligonucleotide
nt: nucleotide(s)
PNA: <u>P</u>eptide <u>N</u>ucleic <u>A</u>cid
IAV: influenza <u>A v</u>irus

## SUPPORTING INFORMATION

**S1 Table. *D. melanogaster* unique transcripts with the potential of forming at least one R12 double-stranded region, identified using *RNAMotif*.**

