## Supplemental Table 1 for "*TFOFinder*: Python program for identifying purine-only double-stranded stretches in the predicted secondary structure(s) of RNA targets"

**S1 Table.** ***D. melanogaster*** **unique transcripts with the potential of forming at least one R12 double-stranded region, identified using *RNAmotif* .**

| Number | Length | Name | FlyBase Gene ID |
|---|---|---|---|
| 1 | 486 | lncRNA:CR44598-RA | FBgn0265809 |
| 2 | 1,297 | lncRNA:CR44619-RA | FBgn0265830 |
| 3 | 1,641 | CR43972-RA | FBgn0264703 |
| 4 | 1,899 | Snp-RD | FBgn0265192 |
| 5 | 2,122 | qkr58E-1-RC | FBgn0022986 |
| 6 | 2,172 | CG9776-RB | FBgn0027866 |
| 7 | 2,306 | CG6294-RB | FBgn0030640 |
| 8 | 2,452 | Rbp1-like-RB | FBgn0030479 |
| 9 | 2,551 | CG6294-RA | FBgn0030640 |
| 10 | 2,551 | CG6299-RB | FBgn0030641 |
| 11 | 2,554 | CG32052-RD | FBgn0044328 |
| 12 | 2,565 | CG32052-RE | FBgn0044328 |
| 13 | 2,903 | Dsp1-RD | FBgn0278608 |
| 14 | 2,996 | repo-RA | FBgn0011701 |
| 15 | 3,643 | eag-RE | FBgn0000535 |
| 16 | 3,649 | eag-RC | FBgn0000535 |
| 17 | 3,652 | CG44422-RF | FBgn0265595 |
| 18 | 3,670 | eag-RF | FBgn0000535 |
| 19 | 3,758 | lncRNA:CR44662-RA | FBgn0265873 |
| 20 | 3,915 | Dsp1-RB | FBgn0278608 |
| 21 | 3,953 | amn-RA | FBgn0086782 |
| 22 | 3,990 | Dsp1-RA | FBgn0278608 |
| 23 | 4,031 | kirre-RC | FBgn0028369 |
| 24 | 4,055 | kirre-RD | FBgn0028369 |
| 25 | 4,139 | Hex-A-RC | FBgn0001186 |
| 26 | 4,159 | CG12075-RD | FBgn0030065 |
| 27 | 4,195 | CG12075-RB | FBgn0030065 |
| 28 | 4,240 | Dsp1-RE | FBgn0278608 |
| 29 | 4,257 | hig-RC | FBgn0010114 |
| 30 | 4,332 | hig-RD | FBgn0010114 |
| 31 | 4,366 | kirre-RE | FBgn0028369 |
| 32 | 4,394 | CG44422-RA | FBgn0265595 |
| 33 | 4,405 | CG8034-RA | FBgn0031011 |
| 34 | 4,471 | mthl1-RA | FBgn0030766 |
| 35 | 4,509 | spoon-RC | FBgn0263987 |
| 36 | 4,511 | Dsp1-RC | FBgn0278608 |
| 37 | 4,525 | Drak-RC | FBgn0052666 |
| 38 | 4,588 | CG15760-RB | FBgn0030508 |
| 39 | 4,754 | lncRNA:CR44754-RA | FBgn0265967 |
| 40 | 4,837 | dpr8-RB | FBgn0052600 |

| | | | |
|---|---|---|---|
| 41 | 4,847 | Drak-RD | FBgn0052666 |
| 42 | 4,953 | kirre-RG | FBgn0028369 |
| 43 | 4,958 | dpr8-RA | FBgn0052600 |
| 44 | 5,083 | mthl1-RB | FBgn0030766 |
| 45 | 5,390 | bi-RE | FBgn0000179 |
| 46 | 5,402 | bi-RD | FBgn0000179 |
| 47 | 5,433 | kirre-RB | FBgn0028369 |
| 48 | 5,496 | RSG7-RB | FBgn0024941 |
| 49 | 5,518 | ovo-RC | FBgn0003028 |
| 50 | 5,533 | CG43736-RG | FBgn0263993 |
| 51 | 5,570 | Hs3st-A-RA | FBgn0053147 |
| 52 | 5,590 | Frq1-RE | FBgn0030897 |
| 53 | 5,594 | Frq1-RD | FBgn0030897 |
| 54 | 5,613 | bi-RG | FBgn0000179 |
| 55 | 5,685 | CG42260-RA | FBgn0259145 |
| 56 | 5,685 | CG42260-RE | FBgn0259145 |
| 57 | 5,686 | Frq1-RB | FBgn0030897 |
| 58 | 5,724 | Hs3st-A-RB | FBgn0053147 |
| 59 | 5,732 | Frq1-RC | FBgn0030897 |
| 60 | 5,815 | CG43736-RF | FBgn0263993 |
| 61 | 5,934 | CG12535-RF | FBgn0029657 |
| 62 | 5,946 | ovo-RA | FBgn0003028 |
| 63 | 5,948 | ovo-RD | FBgn0003028 |
| 64 | 5,977 | kirre-RF | FBgn0028369 |
| 65 | 6,102 | CG42260-RD | FBgn0259145 |
| 66 | 6,606 | ovo-RB | FBgn0003028 |
| 67 | 6,763 | ovo-RE | FBgn0003028 |
| 68 | 6,786 | Sh-RI | FBgn0003380 |
| 69 | 6,816 | CG11000-RC | FBgn0263353 |
| 70 | 6,951 | Zdhhc8-RD | FBgn0085478 |
| 71 | 7,047 | CG14441-RA | FBgn0029895 |
| 72 | 7,244 | BRWD3-RA | FBgn0011785 |
| 73 | 7,264 | CG42260-RB | FBgn0259145 |
| 74 | 7,387 | CG43736-RI | FBgn0263993 |
| 75 | 7,521 | dnc-RR | FBgn0000479 |
| 76 | 7,668 | CG43736-RL | FBgn0263993 |
| 77 | 7,705 | CG44422-RD | FBgn0265595 |
| 78 | 8,094 | B4-RC | FBgn0023407 |
| 79 | 8,223 | br-RA | FBgn0283451 |
| 80 | 8,328 | dnc-RQ | FBgn0000479 |
| 81 | 8,691 | dnc-RT | FBgn0000479 |
| 82 | 9,166 | Myc-RB | FBgn0262656 |
| 83 | 9,215 | eag-RA | FBgn0000535 |
| 84 | 9,455 | eag-RG | FBgn0000535 |
| 85 | 9,456 | dnc-RU | FBgn0000479 |
| 86 | 9,497 | eag-RD | FBgn0000535 |
| 87 | 9,503 | eag-RB | FBgn0000535 |

| | | | |
|---|---|---|---|
| 88 | 9,576 | dnc-RS | FBgn0000479 |
| 89 | 9,717 | Octalpha2R-RC | FBgn0038653 |
| 90 | 9,804 | Octalpha2R-RA | FBgn0038653 |
| 91 | 9,804 | Octalpha2R-RB | FBgn0038653 |
| 92 | 9,822 | br-RL | FBgn0283451 |
| 93 | 10,159 | dnc-RP | FBgn0000479 |
| 94 | 10,706 | lncRNA:CR43314-RD | FBgn0263019 |
| 95 | 11,128 | dnc-RN | FBgn0000479 |
| 96 | 12,088 | brat-RC | FBgn0010300 |
| 97 | 12,181 | Sh-RC | FBgn0003380 |
| 98 | 13,753 | Dora-RB | FBgn0085430 |
| 99 | 14,496 | Smr-RF | FBgn0265523 |
| 100 | 14,745 | rg-RM | FBgn0266098 |
| 101 | 15,201 | Smr-RG | FBgn0265523 |
| 102 | 15,534 | rg-RN | FBgn0266098 |
| 103 | 15,770 | RyR-RB | FBgn0011286 |
| 104 | 15,770 | RyR-RD | FBgn0011286 |
| 105 | 15,788 | RyR-RF | FBgn0011286 |
| 106 | 15,811 | RyR-RJ | FBgn0011286 |
| 107 | 15,812 | RyR-RA | FBgn0011286 |
| 108 | 15,847 | RyR-RC | FBgn0011286 |
| 109 | 15,871 | RyR-RI | FBgn0011286 |
| 110 | 15,981 | rg-RL | FBgn0266098 |
| 111 | 16,003 | RyR-RE | FBgn0011286 |
| 112 | 16,042 | RyR-RH | FBgn0011286 |
| 113 | 16,056 | RyR-RG | FBgn0011286 |
| 114 | 16,076 | CG15465-RA | FBgn0029746 |
| 115 | 16,076 | rg-RR | FBgn0266098 |
| 116 | 16,491 | rg-RQ | FBgn0266098 |
| 117 | 17,173 | CG34417-RW | FBgn0085446 |
| 118 | 17,444 | CG34417-RL | FBgn0085446 |
| 119 | 17,795 | CG34417-RH | FBgn0085446 |
| 120 | 19,353 | CG15465-RB | FBgn0029746 |
| 121 | 19,353 | rg-RP | FBgn0266098 |
| 122 | 20,324 | lncRNA:CR46003-RB | FBgn0267665 |
| 123 | 21,024 | lncRNA:CR46003-RA | FBgn0267665 |
